# Area-specific Biased Global Efficiency in Functional Connectivity Provides Features Negatively Correlated with Age

**DOI:** 10.1101/2020.04.22.054627

**Authors:** Hiroyuki Akama, Airi Ota

## Abstract

It has been acknowledged that graph-theoretical coefficients computed from the adjacency matrix of cerebral resting-state functional connectivity (RSFC) represent aging of the brain and its plasticity facilitating cognitive reserve. In particular, global efficiency (GE) has been recognized as a crucial graph index for age-dependent RSFC. Using the dataset of the Nathan Kline Institute-Rockland Sample [NKI-RS], we found that the regions of the brain in which GE values decay with age were located in the subcortical zone and the cerebellum, whereas an opposite relationship was found in many frontal and parietal regions. Based on this systematic tendency, a new coefficient was proposed that corrects GE, referred to as biased GE (BGE); BGE is calculated by changing the sign of the weight between the superior and the inferior parts of the brain before separately averaging the respective sign groups of area-specific corrected GE values, to influence the raw global network GE. The BGE showed a significant negative correlation with age, irrespective of the scan condition, and strong consistency as an information source of a subject’s identity. We propose that this new index could play an important role in the clinical context of preventive medicine and the maintenance of healthy brains.

## 1. Introduction

In our increasingly ageing society, the burden of dementia has become a global healthcare issue. In light of these circumstances, people interested in preventive medicine have noticed the potential benefits of information provided by studying resting-state functional connectivity (RSFC). This technique is increasingly recognized as a key method to detect the subtle neurocognitive features that characterize the properties and identity (fingerprint) of an individual (Finn et al., 2015). It has been widely recognized, for example, that RSFC can provide information about deterioration associated with aging seen in different aspects of the functional network, including changes in associative weight, dynamic connectivity adjustment, and values of graph theoretical coefficients.

However, compared to other magnetic resonance imaging (MRI)-based information involving T1-weighted imaging or diffusion tensor imaging (DTI), i.e. grey matter volume or fractional anisotropy, respectively, functional MRI data are influenced by scanner surroundings and parameter settings. Therefore, comparing these data across different medical institutions is problematic despite several recent efforts to standardize the criteria and harmonization. MRI datasets recorded from the same subjects who experienced different scanner environments either longitudinally (e.g., the Single Individual volunteer for Multiple Observations across Networks [SIMON] dataset, with a single subject scanned at multiple sites with different scanner models) or inter-sessionally (e.g., the Nathan Kline Institute-Rockland Sample [NKI-RS] dataset, with subjects experiencing three scan conditions with largely different repetition times [TRs]) are valuable for identifying robust markers that characterize individual brains without being affected by measurement conditions.

In addition to the consequence of such datasets, it is noteworthy that several research attempts have partially succeeded in revealing cohesive features from RSFC. In this study, we highlight the importance of graph-theoretical coefficients computed from the adjacency matrix of FC that represents aging of the brain and its plasticity favorable for cognitive reserve. Global efficiency (GE; Latora, 2001) has been recognized as a crucial graph index of brain connectivity (Rubinov and Sporn, 2010; Braun et al., 2011). GE is the average inverse shortest path length over the exhaustive combination of different nodes in a graph. It is acknowledged as a scale to evaluate how efficiently information is transmitted within a network (Latora and Marchiori, 2001). GE can be computed as an index to be assigned to each node, by setting all the other nodes as its targets. Achard and Bullmore (2007) found that in frontal and temporal cortical and subcortical regions, GE was negatively correlated with age.

However, despite the informativeness of age-related GE, proposing a quotient based on whole-brain FC remains an open question, although grey matter volume or fractional anisotropy have been used as sources for computing the Brain Healthcare Quotient (BHQ; Nemoto et al, 2017). In this article, we broaden the scope of monitoring brain health by incorporating functional connectivity into this purpose. Using the cross-sectional dataset of NKI-RS, a new Brain Healthcare Quotient (BHQ) is proposed, accounting for degeneration of brain function through our own corrected area-specific GE. The reason of this adjustment is that the interpretation of GE may vary according to different areas of the brain. Our BHQ, which can be calculated using the IQ-like formula: 100 + 15 × (concerned coefficient − mean) / standard deviation, can be named the Functional Connectivity (FC)-BHQ given the previous quotients, Grey-Matter (GM)- BHQ and Fractional Anisotropy (FA)-BHQ (Nemoto et al., 2017).

## 2. Materials and Methods

We used the dataset of Nathan Kline Institute Rockland Sample NKI RS Release 1 (http://fcon_1000.projects.nitrc.org/indi/pro/nki.html) in which the following three scan protocols were adopted for acquiring resting state functional MRI (RS-fMRI) volumes on a SIEMENS MAGNETOM TrioTim syngo 3.0T scanner (Siemens Healthineers AG, Erlangen, Germany): i) TR=645 ms, echo time [TE]=30 ms, multiband factor= 4, slice numbers=40, FOV=222×222 mm^2^, slice thickness=3 mm, flip angle= 80°, voxel size= 3.0×3.0×3.0 mm, duration time=10 minutes; ii) TR=1400 ms, TE=30 ms, multiband factor=4, slice numbers=64, FOV=224×224 mm^2^, slice thickness=2 mm, flip angle = 65°, voxel size = 2.0×2.0×2.0 mm, duration time=10 minutes; and iii) TR=2500 ms, TE=30 ms, slice numbers=38, FOV=240×212 mm^2^, slice thickness=3.2 mm, flip angle=80°, voxel size=3.0×3.0×3.0 mm, duration time=5 minutes. We refer to the sessions i), ii) and iii) as 645 ms, 1400 ms, and 2500 ms, respectively. Sixty healthy volunteers who experienced these three sessions were examined in the resting state fMRI analysis. They were 34 females and 26 males, of whom 56 were right-handed and 4 were left-handed, and were aged between 10 and 83 years with a mean age of 39.2 years.

FC analyses were performed using the CONN toolbox v18a (https://web.conn-toolbox.org/), with the default frequency band (<0.10 Hz). Irrespective of the scan condition (different TRs), we resliced with the isotropic voxel size of 2.0×2.0×2.0 mm and smoothed, with an 8-mm Gaussian kernel, all the functional volumes normalized by the standard template of the Montreal Neurological Institute (MNI). After the second level analysis, a table of graph theoretical coefficients (degree, cost, local efficiency, GE, betweenness centrality, and clustering coefficient) was automatically generated for each subject and each area of the CONN brain atlas, named “atlas.nii” and “networks.nii.” These NIFTI files stemmed from the FSL Harvard-Oxford Atlas, as well as the cerebellar areas from the Automated Anatomical Labelling (AAL) Atlas (Whitfield-Gabrieli & Nieto-Castanon, 2012). This subject/area-specific graph theoretical table includes 132 regions of interest (ROIs) and 32 representative nodes of the eight intrinsic networks (default mode, sensorimotor, visual, saliency, dorsal attention, frontoparietal, language, and cerebellar), so a geodesic redundancy was built in by the partial duplication of areas shared by the two NIFTI maps. However, we left this duplication untouched, only to enhance a clear understanding of functional topography. In extracting the GE information from the graph theoretical tables corresponding to the TRs of 645 ms, 1400 ms, and 2500 ms, a correlation analysis was carried out across the three mean GE values (of the whole brain networks) for individual subjects, and between age and area-specific GE for each within-scan condition.

This dataset is shared and open to the public through NITRC (NeuroImaging Tools & Resources Collaboratory), and we performed this study under the approval of the Institutional Review Board of Tokyo Institute of Technology, Japan (authorization number: A19106). Regarding statistical analysis, MATLAB 2018b was used for creating scripts to compute the FC-BHQ from the result files of the CONN toolbox. In this study, the statistical threshold was set at p < 0.05 for the correlation test.

## 3. Results

When analyzing the average GE values for the three sessions that each subject underwent with the different scan parameters, a pairwise-correlation was found to be statistically significant between the TRs of 1400 ms and 2500 ms (r=0.4674, p=0.0002). However, between 645 ms and 1400 ms (r=0.2265), and between 645 ms and 2500 ms (r=0.1073), the correlations were not significant. Regarding the within-scan condition analysis, the GE of the whole brain functional networks significantly decreased with age under the scan condition of TR=1400 ms (r=−0.3305, p=0.01), while the results for the other conditions were r=0.085 (n.s.) for 645 ms and r=−0.181 (n.s.) for 2500 ms.

When examining the correlation between the GE of each area as a node and the years of subjects, some areas were found to elicit significant effects of aging (|r|>0.25, p<0.05). Table 1, divided by rows for the different TR sessions, indicates the area-nodes where the GE negatively correlates with age, using deep green (p<0.01) and light green (p<0.05) background colors to indicate the degree of statistical significance. These area-nodes showing traces of senile change were depicted by cool colors in the left three rendered brain images of Figure 1, together with those that showed significantly positive correlations, which were depicted by warm colors. The rightmost image of Figure 1 displays mapping of all the 164 area-nodes by a heat map scale according to the sum of ranks for the GE-age correlation across the three sessions. The top three area-nodes for this ranking, which recorded the most negative correlation between GE and age, were the left Temporal Fusiform Cortex Posterior Division (rank sum = 7), the left Inferior Temporal Gyrus Temporooccipital part (rank sum = 24), and the Vermis 6 (rank sum = 30). Conversely, the bottom three, where GE increased with aging to the maximal degree, were the Cingulate Gyrus posterior division (rank sum = 477), the Supramarginal Gyrus in Salience network (rank sum = 463), and the Left Superior Parietal Lobule (rank sum = 443).

**Table 1.** Anatomical areas in which the GE correlated negatively with age in each of the three scanning conditions (Top: repetition time [TR]=645 ms; Middle: TR=1400 ms; Bottom: TR=2500 ms, Background colors: deep green (p<0.01) and light green (p<0.05)).

| 645 |  |  | 301 |
| --- | --- | --- | --- |
| rank | regions | Correlation coefficient | p-value |
| 1 | Vermis45 | -0.3554 | 0.0053 |
| 2 | ParahippocampalGyrus AnteriorDivision L | -0.3541 | 0.0055 |
| 3 | TemporalFusiformCortex PosteriorDivision L | -0.3382 | 0.0082 |
| 4 | InferiorTemporalGyrus TemporooccipitalPart L | -0.3369 | 0.0085 |
| 5 | TemporalFusiformCortex PosteriorDivision R | -0.3310 | 0.0098 |
| 6 | Vermis6 | -0.3173 | 0.0135 |
| 7 | Cerebelum45 L | -0.3114 | 0.0154 |
| 8 | ParahippocampalGyrus AnteriorDivision R | -0.2975 | 0.0210 |
| 9 | ParahippocampalGyrus PosteriorDivision R | -0.2853 | 0.0271 |
| 10 | Caudate L | -0.2745 | 0.0338 |
| 11 | Vermis3 | -0.2743 | 0.0339 |
| 12 | TemporalFusiformCortex AnteriorDivision R | -0.2728 | 0.0350 |
| 13 | Vermis7 | -0.2707 | 0.0364 |
| 14 | Caudate R | -0.2603 | 0.0446 |

| 1400 |  |  |  |
| --- | --- | --- | --- |
| rank | regions | Correlation coefficient | *p*-value |
| 1 | **CaudateR** | **-0.4884** | **0.0001** |
| 2 | **CaudateL** | **-0.4584** | **0.0002** |
| 3 | **TemporalFusiformCortex PosteriorDivision L** | **-0.4213** | **0.0008** |
| 4 | **Accumbens L** | **-0.4070** | **0.0013** |
| 5 | **Cerebelum8 R** | **-0.3877** | **0.0022** |
| 6 | **SubcallosalCortex** | **-0.3650** | **0.0041** |
| 7 | **Amygdala L** | **-0.3634** | **0.0043** |
| 8 | **Accumbens R** | **-0.3528** | **0.0057** |
| 9 | **Cerebelum6 R** | **-0.3456** | **0.0068** |
| 10 | **InferiorFrontalGyrus ParsOpercularis L** | **-0.3415** | **0.0076** |
| 11 | **ParahippocampalGyrus PosteriorDivision L** | **-0.3378** | **0.0083** |
| 12 | Cerebelum10 L | -0.3295 | 0.0101 |
| 13 | Vermis6 | -0.3210 | 0.0124 |
| 14 | OccipitalFusiformGyrus R | -0.3164 | 0.0138 |
| 15 | TemporalFusiformCortex PosteriorDivision R | -0.3163 | 0.0138 |
| 16 | InferiorTemporalGyrus _TemporooccipitalPart L | -0.3088 | 0.0164 |
| 17 | MiddleTemporalGyrus TemporooccipitalPart L | -0.3039 | 0.0182 |
| 18 | Vermis45 | -0.2975 | 0.0210 |
| 19 | Cerebelum10 R | -0.2951 | 0.0221 |
| 20 | MiddleTemporalGyrus AnteriorDivision R | -0.2950 | 0.0221 |
| 21 | Amygdala R | -0.2899 | 0.0247 |
| 22 | Cerebelum3 L | -0.2863 | 0.0266 |
| 23 | ParahippocampalGyrus PosteriorDivision R | -0.2857 | 0.0269 |
| 24 | Thalamus L | -0.2857 | 0.0269 |
| 25 | InferiorTemporalGyrus PosteriorDivision R | -0.2854 | 0.0271 |
| 26 | MiddleTemporalGyrus AnteriorDivision L | -0.2685 | 0.0380 |
| 27 | FrontalMedialCortex | -0.2660 | 0.0399 |
| 28 | MiddleTemporalGyrus PosteriorDivision R | -0.2652 | 0.0406 |

| 2500 |  |  |  |
| --- | --- | --- | --- |
| rank | regions | Correlation coefficient | *p*-value |
| 1 | TemporalFusiformCortex PosteriorDivision L | -0.4294 | 0.0006 |
| 2 | Cerebelum6 R | -0.4134 | 0.0010 |
| 3 | SuperiorTemporalGyrus AnteriorDivision R | -0.3620 | 0.0045 |
| 4 | InferiorTemporalGyrus TemporooccipitalPart L | -0.3603 | 0.0047 |
| 5 | CerebelumCrus1 R | -0.3538 | 0.0056 |
| 6 | Cerebelum45 R | -0.3388 | 0.0081 |
| 7 | ParahippocampalGyrus PosteriorDivision L | -0.3214 | 0.0123 |
| 8 | LateralOccipitalCortex InferiorDivision R | -0.3194 | 0.0129 |
| 9 | Cerebelum45 L | -0.3191 | 0.0130 |
| 10 | TemporalPole L | -0.3126 | 0.0150 |
| 11 | Vermis6 | -0.3112 | 0.0155 |
| 12 | Cerebelum8 R | -0.3110 | 0.0156 |
| 13 | TemporalFusiformCortex PosteriorDivision R | -0.3081 | 0.0166 |
| 14 | ParahippocampalGyrus PosteriorDivision R | -0.3000 | 0.0199 |
| 15 | SuperiorTemporalGyrus AnteriorDivision L | -0.2987 | 0.0204 |
| 16 | Caudate R | -0.2973 | 0.0211 |
| 17 | InferiorTemporalGyrus TemporooccipitalPart R | -0.2860 | 0.0268 |
| 18 | Cerebelum10 L | -0.2843 | 0.0277 |
| 19 | Cerebelum8 L | -0.2789 | 0.0310 |
| 20 | MiddleTemporalGyrus AnteriorDivision R | -0.2652 | 0.0406 |
| 21 | TemporalOccipitalFusiformCortex L | -0.2568 | 0.0476 |
| 22 | Vermis8 | -0.2546 | 0.0496 |

**Figure 1:**
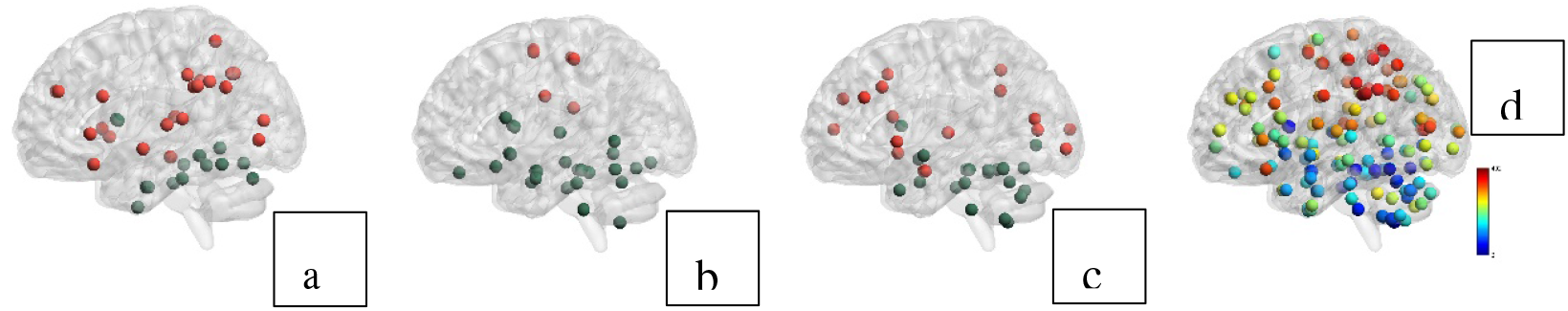
Correlation patterns between global efficiency (GE) and age. The rendered brain images a, b, and c show the area nodes where the GEs were positively (warm colors in the superior part of the brain) and negatively (cool colors in the inferior part of the brain) correlated with age (a: repetition time [TR]=645 ms; b: TR=1400 ms; and c: TR=2500 ms). The image d displays the mapping of all the 164 area-nodes by a heat map scale according to the sum of ranks for the GE-age correlation across the three sessions.

These results convincingly revealed that the regions in which GE values decay with age are mostly located in the subcortical zone and the cerebellum, that is, the inferior part of the brain. Conversely, the GE was elevated in the superior part of the brain, particularly in the parietal lobe. This intriguing global distribution pattern enabled us to define an area-specific binary weight vector to correct the whole brain GE for each participant. We assigned the value of plus one to every node of the frontal, parietal and occipital lobes, and minus one to each of the remaining regions, comprising the temporal lobe, basal ganglia, limbic system, and cerebellum. After generating the inner product of the weight vector and the list of the area-specific GE values for each participant, we partitioned the elements into two subsets according to their positive and negative signs, separately computed the means, and summated them to produce the area bias to be added to each participant’s global network GE (average GE across areas). This correction of GE plus area bias determined the biased global efficiency (BGE).

BGE was effective in preserving the connectivity features of the individual participants, as well as expressing alteration and degeneration of the brain functionality due to the effects of increasing age. The pairwise correlations across the participants’ BGE records were all highly significant between any two combinations from the three scan sessions: between 1400 ms and 2500 ms (r=0.8354, p=1.04e-16), between 645 ms and 1400 ms (r=0.9091, p=9.97e-24), and between 645 ms and 2500 ms (r=0.7556, p=3.17e-12). In the within-scan condition analysis, the BGE of the whole brain functional networks significantly decreased with age, with correlation values smaller than −0.3 in all scan conditions: TR=645 ms (r= −0.3027, p=0.019), 1400 ms (r= −0.3728, p=0.003), and 2500 ms (r= −0.3211, p=0.012; Figure 2 Left). The linear regression analysis was significant between the age and BGE means across the three scans. The right part of Figure 2 represents the scatter plot and regression line of age versus FC-BHQ, calculated as shown in the Introduction.

**Figure 2:**
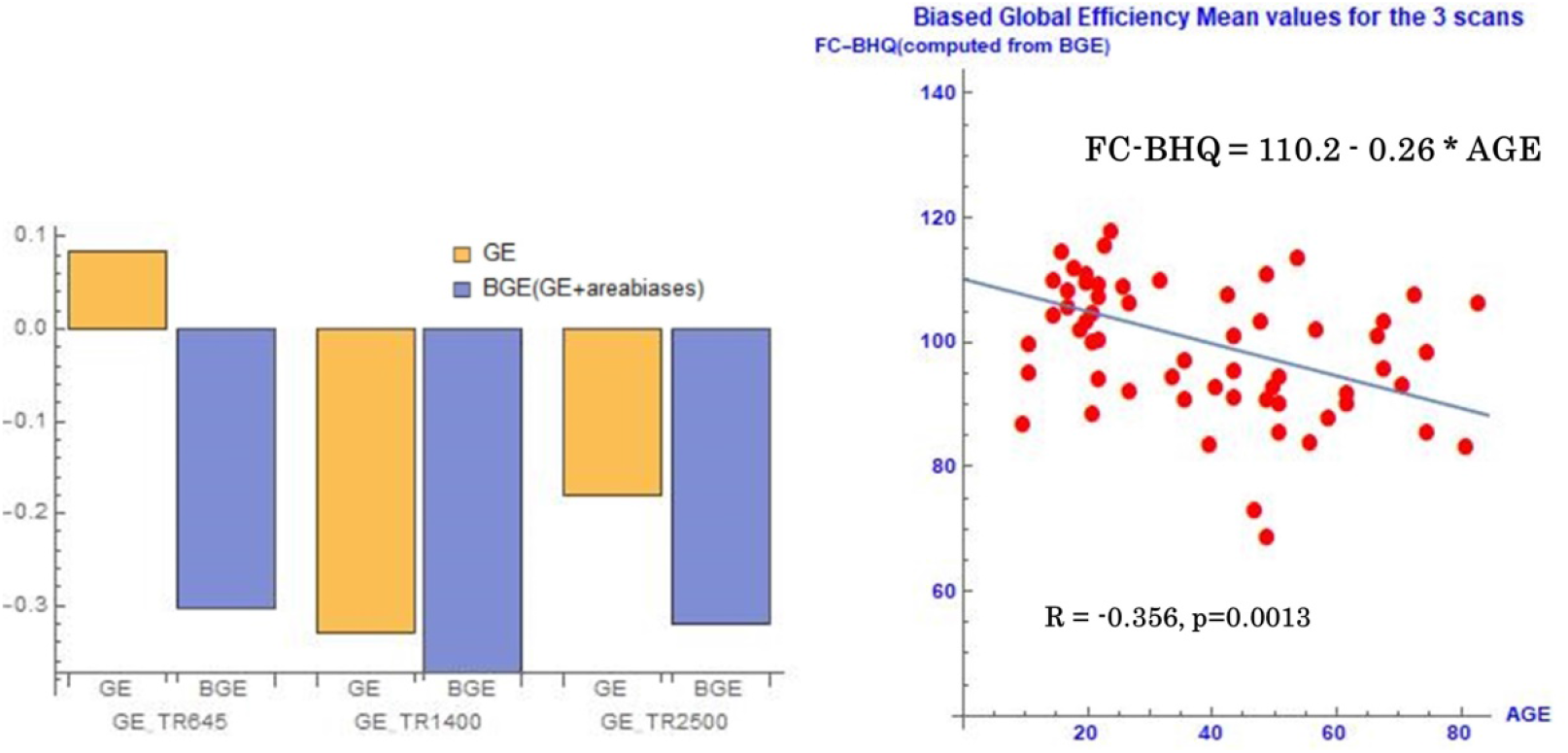
The effectiveness of biased global efficiency (BGE). Left: The BGE, by adding area biases, improved the global efficiency (GE) of the whole network of RSFC as an index for age-related information. The correlation between the graph theoretical coefficient and age became statistically negative and stable, irrespective of the scan condition. Right: Scatter plot and the regression line of age versus Functional Connectivity - Brain Healthcare Quotient (FC-BHQ) calculated as: 100 + 15 × (BGE − mean) / standard deviation.

## 4. Discussion

The robust significance of the BGE suggests that this new coefficient is a good candidate for FC-BHQ and is at least comparable to the already-proposed FA-BHQ (Nemoto et al., 2017), in which the reported R was −0.417. Although further refinement will enable more accurate representation of the age-related modulation of FC among individual subjects, the BGE provides important descriptions about the patterns of regional differences and the implications of GE, in particular, information transferability. Interestingly, our findings differ from the results of a previous study evaluating the correlation between area-specific GE and ageing. Achard and Bullmore (2007) suggested that GE significantly decreased through ageing in frontal and temporal cortical and subcortical regions. The dorsal cingulate and middle frontal gyri were highlighted as particular loci of negative correlations. Nevertheless, the NKI-RS fMRI dataset does not corroborate that geodesic pattern in functionality. The majority of the frontal regions in our study elicit non-significant or even significantly positive correlations with age. The right Frontal Operculum (r= 0.3449, p=0.007) and the Cingulate Gyrus posterior division (r= 0.3604, p=0.005) in the 645 ms condition, and the right Inferior Frontal Gyrus pars opercularis (r=0.3698, p=0.004) in the 2500 ms condition, were positively correlated with age. There were only two frontal areas negatively correlated with age, and both were identified in the scan with the TR of 1400 ms, namely the left Inferior Frontal Gyrus pars opercularis (r=−0.3415, rank=10^th^) and the Medial Frontal cortex (r=−0.266, rank=27th). Interestingly, the rank numbers of these areas in the GE list considerably fluctuated according to the TR conditions: in the 645 ms and 2500 ms sessions, ranks of 116^th^ and 144^th^ for the former and 111^th^ and 37^th^ for the latter were observed, respectively.

Thus, the effect of aging appears to influence GE of the superior and inferior parts of the brain differently. The frontal regions may not have elicited connectivity degeneration because of cognitive reserve (CR; Reuter-Lorenz et al., 2008; Franzmeier et al., 2018; Martínez et al., 2017; Colangeli et al., 2016). According to Benson et al. (2018), “higher functional connectivity in fronto-parietal and salience networks may protect against detrimental effects of white matter lesions on executive functions.” The nodal regions associated with higher cognitive functions may be protected against cerebrovascular pathology by the intrinsic networks encompassing the frontal and parietal lobes. This account of the neural basis of cognitive reserve is in line with the enhanced activity of task control networks that was assessed through the analysis of NKI-RS. In fact, there is no area representative of the salience network that recorded a negative correlation with age; instead, there were positive correlations with age in our study, especially in the 2500 ms TR condition, where three areas of the salience network showed a significantly positive correlation with age. These areas were the right Prefrontal cortex with the center at the MNI coordinates of [32 46 27] (r=0.2586, p=0.046), the right Anterior Insula including [47 14 0] (r=0.2838, p=0.028), and the left Supramarginal gyrus including [60 39 31] (r=0.2884, p=0.025).

In contrast, the decline in GE recorded at the temporal cortical and subcortical regions has been attributed to white matter lesions, such as those caused by multiple sclerosis in the elderly (Bullmore et al., 2011; He et al., 2009). Since this coefficient represents functional integration through the shortest path networks, a drop in the GE values of these regions might reflect the emergence of inefficient path routes as a result of multiple grey matter nodes in the functional connectivity graph. These additional routes, and their effect on reducing GE, may result from cerebral calcification or microinfarction, which are averted in order to complement the lost efficient paths. It is well known that the cerebellum, the caudate and the parahippocampal gyrus, where the largest negative correlation between GE and age was observed, are selectively engaged with motion and memory, and show the effects of increasing age. It is our hypothesis that the systematic lowering of GE in the inferior part of the brain follows the routes of such compensation.

The effectiveness of the BGE is based on the cognitive reserve and the neural compensation, which can be contrasted with each other by changing the sign of the weight between the superior and the inferior parts of the brain. However, besides the merits of the BGE as an index of brain aging, it also has another role in the identification of individual brains. From the viewpoint of harmonization across different scan protocols enabling multi-environmental data evaluation in the clinical context, the BGE values listed for subjects exhibited strong robustness for all the TR conditions. The within-subject correlations were 0.8354 between the TRs of 645 ms and 1400 ms, 0.9091 between the TRs of 1400 ms and 2500 ms, and 0.7556 between the TRs of 645 ms and 2500 ms. These results demonstrate that the coefficient can play the role of a “fingerprint,” identifying subjects irrespective of their age, so long as the RSFC data are taken at the same life stage.

We also focused on the anatomical regions where there is a large difference in GE values, dependent on the scan conditions. Accordingly, we ordered the sum of squares of the difference in the ranks of the GE values that each region was associated with in the three TR sessions. The top six regions where the difference in the GE was largest across the scan conditions were the bilateral IFG Pars Opercularis, the right Middle Temporal Gyrus, the right Temporal Pole, and the right Middle Frontal Gyrus. Interestingly, several regions in this list are interconnected by the Extreme Capsule (Saur et al., 2014) that constitutes the ventral pathway of language processing. It remains to be seen whether this phenomenon is reproducible by the other datasets, or whether it is due to fluctuations in cognitive process related to, for example, mind wandering or physical noises associated with scan protocols such as the background scanner sounds produced by echo planar imaging (EPI).

This study has some limitations, some of which may warrant further study. The concept of efficiency may take the value of zero for isolated vertices setting infinity for their shortest paths to the others, and close to zero for the extremities of dangling chains or the nodes within a disconnected subgraph. We found that some subcortical regions tended to record such outliers, which should be removed from area-wise regression modelling between GE or BGE and age. For example, under the TR = 1400 ms condition, there were 8 and 7 subjects whose GE of the right and left caudate nuclei were significantly below the double standard deviation from the mean (0.0318 and 0.055), respectively. The reason for these outliers remains to be delineated and the treatment method should be developed further beyond the present concepts. Furthermore, open questions remain with respect to how we will be able to establish a new method of embedding the BGE into the profile of RSFC for accurately identifying individual subjects. Graph convolutional technology in deep learning (Ktena et al., 2018) would be promising for this purpose; however, our final goal will be finding “fingerprints” of a subject at his/her precise age since the profile itself might be constantly changing by aging.

## 5. Conclusion

Through use of the RSFC dataset from the NKI-RS study, we have shown that GE values decay with age in the subcortical zone and the cerebellum, but in many frontal and parietal regions an opposite pattern was found. Based on our results a corrected GE is proposed, which involves changing the sign of the weight between the superior and the inferior parts of the brain before separately averaging the respective sign groups of area-specific corrected GE values, to influence the raw global network GE. The BGE, as a customized graph coefficient for evaluating age-related systematic rewiring in RSFC, can be utilized not only as a brain healthcare quotient but also as a fingerprint of an individual brain at a particular age. The robustness of the patterns shown through different data samples could facilitate a new clinical research field in neuro-medicine that focuses on the maintenance of brain health. A longitudinal study may become possible by taking advantage of the tractability of this index in pursuing synchronic identifiability as well as diachronic modulation of individual RSFC features.

## 7. Author Contributions

Conceptualization: Hiroyuki Akama and Airi Ota / Data curation and analysis: Hiroyuki Akama and Airi Ota / Investigation: Hiroyuki Akama and Airi Ota / Methodology: Hiroyuki Akama and Airi Ota / Supervision: Hiroyuki Akama / Writing: Hiroyuki Akama and Airi Ota

## Abbreviations

BGE: biased global efficiency
BHQ: brain healthcare quotient
GE: global efficiency
RSFC: resting-state functional connectivity.

## 9. Acknowledgments

The authors would like to express their gratitude to Dr. Yoshinori Yamakawa, ImPACT Program of Council for Science, Technology and Innovation (Cabinet Office, Government of Japan) for important advice.

## Data Availability Statement

The datasets generated for this study are available on request to the corresponding author.

